# Undocumented potential for primary productivity in a globally-distributed bacterial photoautotroph

**DOI:** 10.1101/140715

**Authors:** E.D. Graham, J.F. Heidelberg, B.J. Tully

## Abstract

Aerobic anoxygenic phototrophs (AAnPs) are common in the global oceans and are associated with photoheterotrophic activity. To date, AAnPs have not been identified in the surface ocean that possess the potential for carbon fixation. Using the *Tara Oceans* metagenomic dataset, we have reconstructed draft genomes of four bacteria that possess the genomic potential for anoxygenic phototrophy, carbon fixation via the Calvin-Benson-Bassham cycle, and the oxidation of sulfite and thiosulfate. Forming a monophyletic clade within the *Alphaproteobacteria* and lacking cultured representatives, the organisms compose minor constituents of local microbial communities (0.1-1.0%), but are globally distributed, present in multiple samples from the North Pacific, Mediterranean Sea, the East Africa Coastal Province, and the South Atlantic. These organisms represent a shift in our understanding of microbially-mediated photoautotrophy in the global oceans and provide a previously undiscovered route of primary productivity.

**Significance Statement:** In examining the genomic content of organisms collected during the *Tara Oceans* expedition, we have identified a novel clade within the *Alphaproteobacteria* that has the potential for photoautotrophy. Based on genome observations, these organisms have the potential to couple inorganic sulfur compounds as electron donors to fix carbon into biomass. They are globally distributed, present in samples from the North Pacific, Mediterranean Sea, East Africa Coastal Current, and the South Atlantic. This discovery may require re-examination of the microbial communities in the global ocean to understand and constrain the impacts of this group of organisms on the global carbon cycle.

## Introduction

It has been understood for decades that the basis of the global marine carbon cycle are oxygenic photoautotrophs that perform photoautotrophic processes. Two additional phototrophic processes are common in the ocean and are mediated by proteorhodopsin-containing microorganisms and aerobic anoxygenic phototrophs (AAnPs). Proteorhodopsins and AAnPs have historically been associated with photoheterotrophy^1,2^, a process that supplements additional energy to microorganisms beyond what is obtained as part of a heterotrophic metabolic strategy. AAnPs utilize type-II photochemical reaction centers (RCIIs) and bacteriochlorophyll (BChl), are globally distributed^3^, and have been identified in phylogenetically diverse groups of microorganisms^4-6^. Though anaerobic microorganisms with RCIIs and BChl are known to fix CO_2_^7^ and marine AAnPs can incorporate inorganic carbon via anaplerotic reactions^8^, marine AAnPs have not been linked to carbon fixation^9^. The identification of marine AAnPs capable of carbon fixation adds to our understanding of microbial photosynthesis in the global oceans and represents a previously undiscovered route of photoautotrophy.

The *Tara Oceans* expedition generated microbial metagenomes during a circumnavigation of the global oceans^10,11^. *Tara Oceans* samples were collected from 63 sites in 10 major ocean provinces, with most sites contributing multiple sampling depths (generally, surface, deep chlorophyll maximum [DCM], and mesopelagic) and multiple size fractions (generally, ‘viral’, ‘girus’ [giant virus], ‘bacterial’, and ‘protistan’) from each depth (Supplementary Data File 1). We independently assembled each sample and assemblies from all samples within a province were combined and subjected to binning techniques to reconstruct microbial genomes (Fig. 1 and Extended Data Fig. 1). Microbial genomes reconstructed from eight of ten provinces (Mediterranean, Red Sea, Arabian Sea, Indian Monsoon Gyre, East Africa Coastal, South Atlantic, and North Pacific; 36 sites, 134 samples) were annotated using the KEGG Ontology (KO) system^12^ and examined for the genes and pathways of interest.

**Fig. 1.**
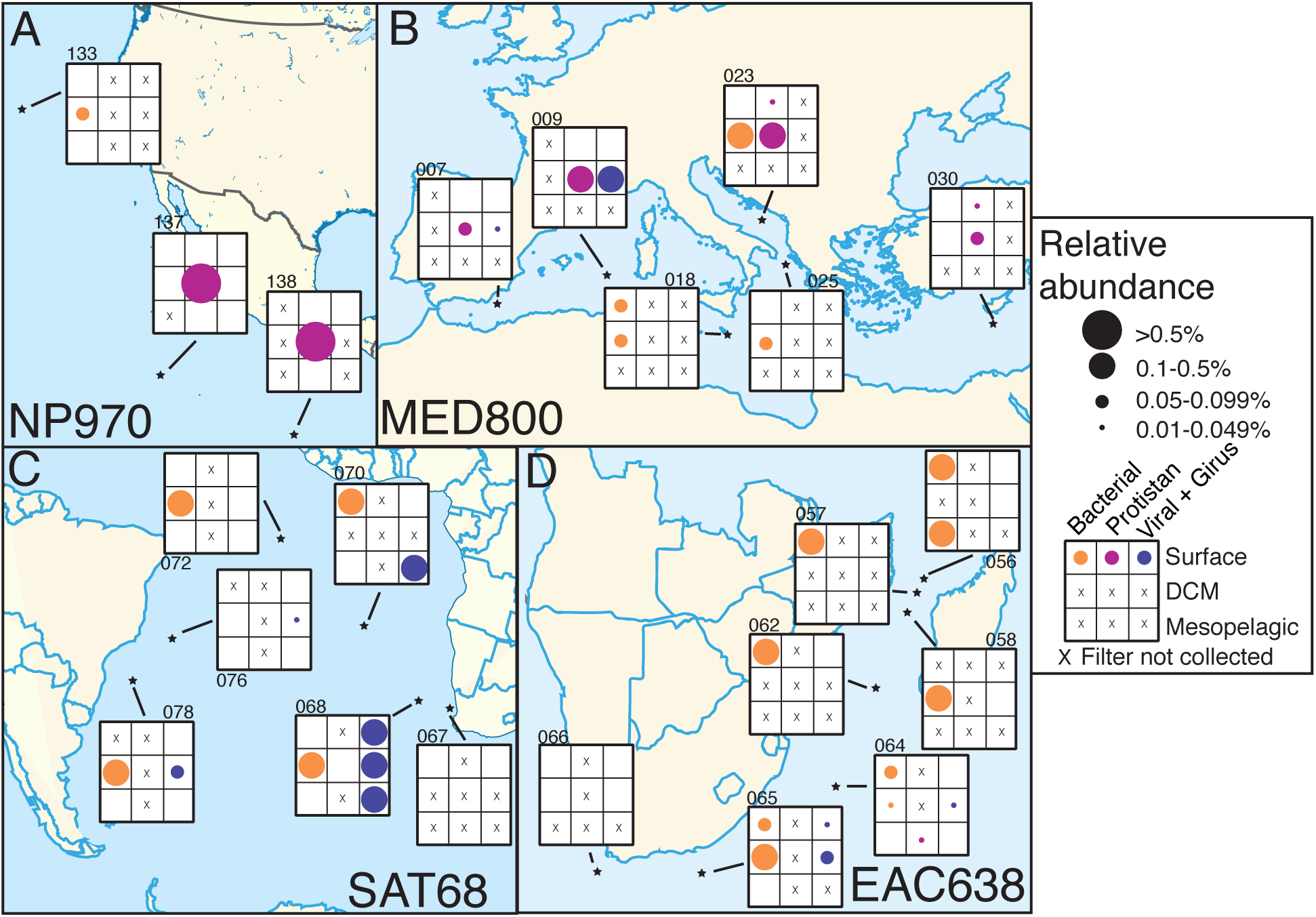
The approximate locations of *Tara Oceans* sampling sites used to generate metagenomes incorporated in to this study. Each grid represents the three possible sample depths and filter fractions (top row: surface, middle row: DCM, bottom row: mesopelagic). An ‘X’ denotes that no sample was collected for that depth and size fraction at the site. Circle size represents relative abundance. (A) North Pacific – NP 970, (B) Mediterranean Sea – MED800, (C) South Atlantic – SAT68, and (D) East Africa current – EAC638. Size fraction: orange, ‘bacterial’ (0.22-1.6μm); purple, ‘protistan’ (0.8-5.0μm); blue, ‘girus’+‘viral’ (<0.22-0.8μm). The maps in Figure 1 were modified under a CC BY-SA 3.0 license from ‘South Atlantic Ocean laea location map’ by Tentotwo, ‘North America laea location map’ by TUBS, ‘Location of Mozambique in Africa’ by Rei-artur, and ‘Blank Map of South Europe and North Africa’ by historicair.

## Results and Discussion

From 1,774 metagenome-assembled genomes (MAGs), 53 genomes possessed the genes encoding the core subunits of RCIIs (PufLM). Of those 53, four genomes (MED800, EAC638, SAT68, NP970; Fig. 1) also contained genes for ribulose-1,5-bisphosphate carboxylase (Rubisco; RbsLS; Fig. 2). Rubisco has four major forms, of which three (Types I, II, and III) have been shown to fix CO_2_ and two are known to participate in the Calvin-Benson-Bassham (CBB) cycle (Types I and II^13,14^. Phylogenetic placement of the Rubisco large subunits recovered from the genomes revealed them to be of the Type IC/D subgroup^13^, suggesting that the identified proteins represent *bona fide* Rubiscos capable of carbon fixation (Fig. 2). Within the Type IC/D subgroup, the Rubisco sequences from the four analyzed genomes formed a distinct cluster with environmental sequences derived from the Global Ocean Survey (GOS) metagenomes^15,16^, but lacking sequences from reference organisms.

**Fig. 2.**
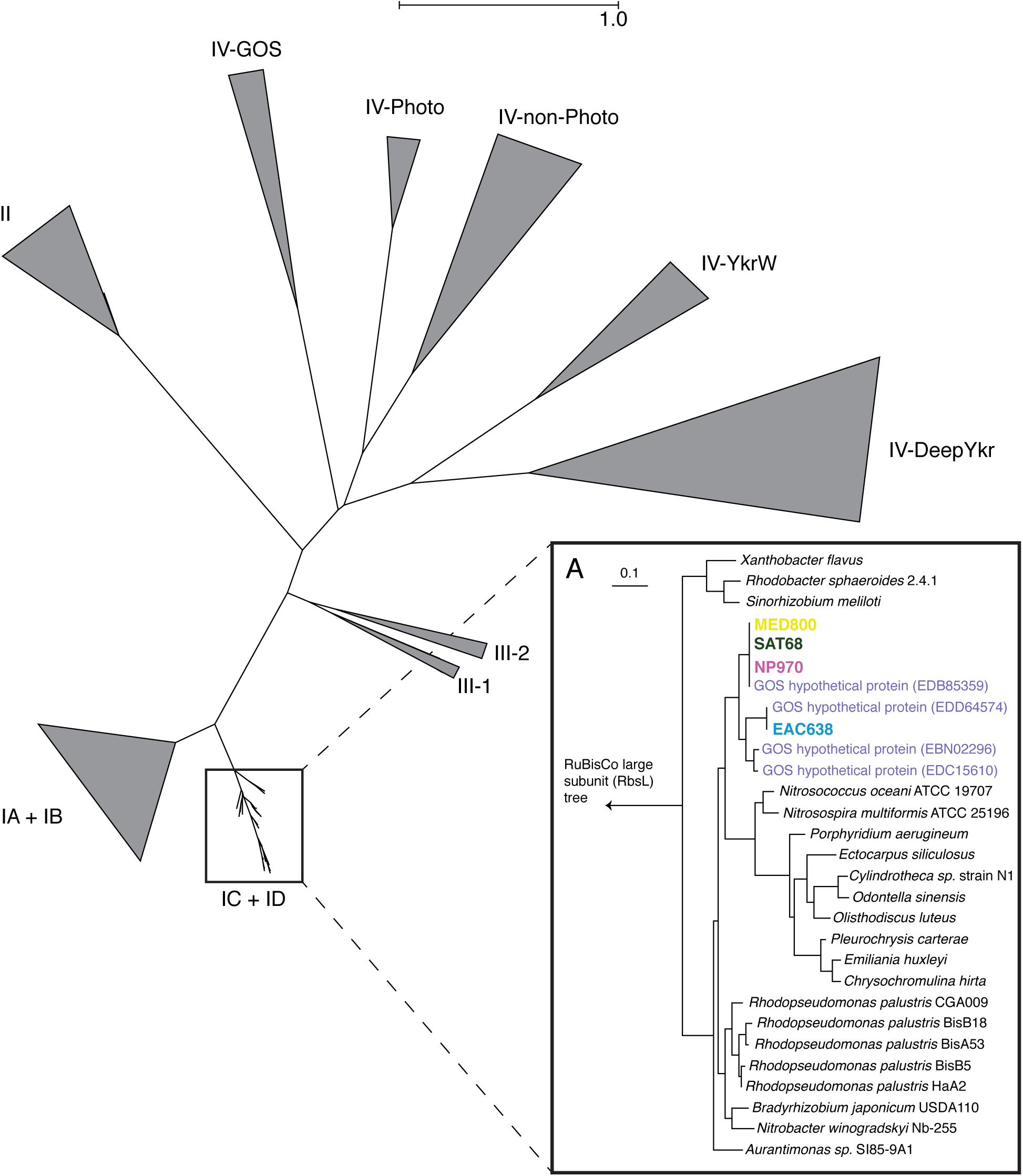
Phylogenetic tree of the ribulose-1,5-bisphosphate carboxylase large subunit (Rubsico, RbsL) with the major forms denoted. Inset (**A**) – A zoomed in view of the Type 1C/D Rubsico subgroup. Purple sequence names denote RbsL proteins from the Global Ocean Survey. Sequences used for this tree can be found in Supplementary Data File 10. Phylogenetic distances and local support values can be found in Supplementary Data File 12. Sequence information, including accession numbers and assignments can be found in Supplementary Data File 8.

Similarly, the PufM sequences from the analyzed *Tara* genomes did not cluster with reference sequences, instead grouping with sequences from the GOS metagenomes^15^. Sequences from MED800, SAT68, and NP970 were group together in one cluster, while EAC638 was located in a separate cluster (Fig. 3). The MED800/SAT68/NP970 clade is basal to the previously identified phylogroups E and F, while the EAC638 clade is basal to the *Roseobacter*-related phylogroup G^17^. As MED800, EAC638, SAT68, and NP970 branch in distinct clades on both the RbsL and PufM trees that consist of entirely environmental sequences, it may be possible that these clades represent a phylogenetically coherent group of organisms with the potential for both phototrophy and carbon fixation.

**Extended Data Fig. 3.**
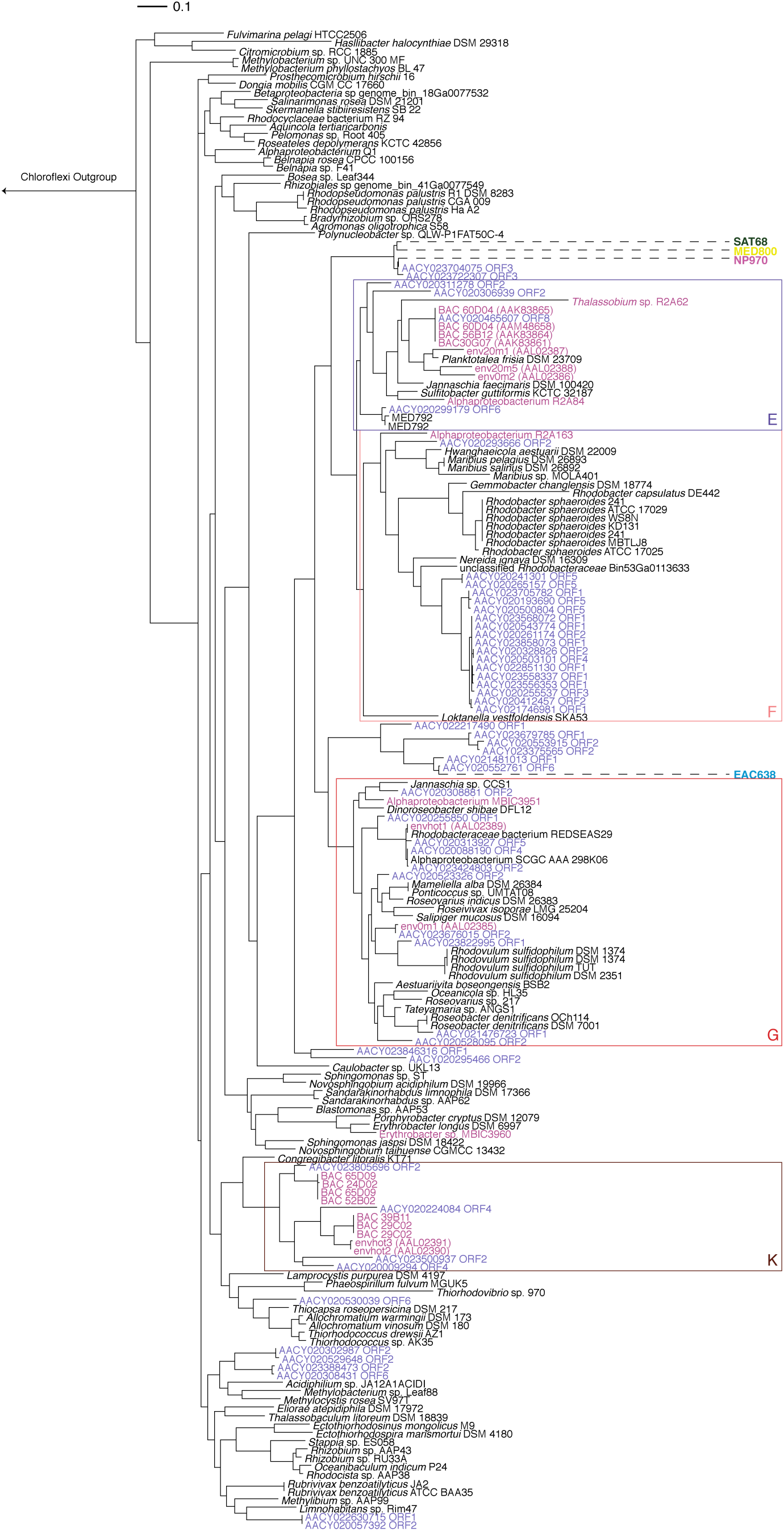
Phylogenetic tree of the M subunit of type-II photochemical reaction center (PufM). Environmental sequences obtained from the Global Ocean Survey (purple, •) and Béjá *et al.* (2002) (pink, •) are highlighted. Boxes illustrate approximate positions of phylogroups previously assigned by Yutin *et al.* (2007). Sequences used for this tree can be found in Supplementary Data File 11. Phylogenetic distances and local support values can be found in Supplementary Data File 13. Sequence information, including accession numbers and taxonomies can be found in Supplementary Data File 9.

The draft genomes were of high enough quality (66-85% complete; <5.5% duplication; Table 1) to possess sufficient phylogenetic markers for accurate placement (Extended Data Table 1). The four organisms form a monophyletic clade basal to the Family *Rhodobacteraceae* (Fig. 4). The relationship between the genomes would suggest that NP970, SAT68, and MED800 are phylogenetically more closely related to each other than either are to EAC638. As is common with assembled metagenomic sequences, the recovered genomes lack a distinguishable 16S rRNA gene sequence. However, based on the observed phylogenetic distance in the concatenated marker tree, we suggest that these organisms represent a new clade within the *Rhodobacteraceae*, and possibly a family-level clade previously without a reference sequence within the *Alphaproteobacteria*. We propose that NP970, SAT68, and MED800 represent three species within the same genus (tentatively named, ‘*Candidatus Luxescamonas taraoceani’*), with EAC638 as a representative of a species in a sister genus (tentatively named, ‘*Candidatus Luxescabacter africus’*).

**Table 1.**
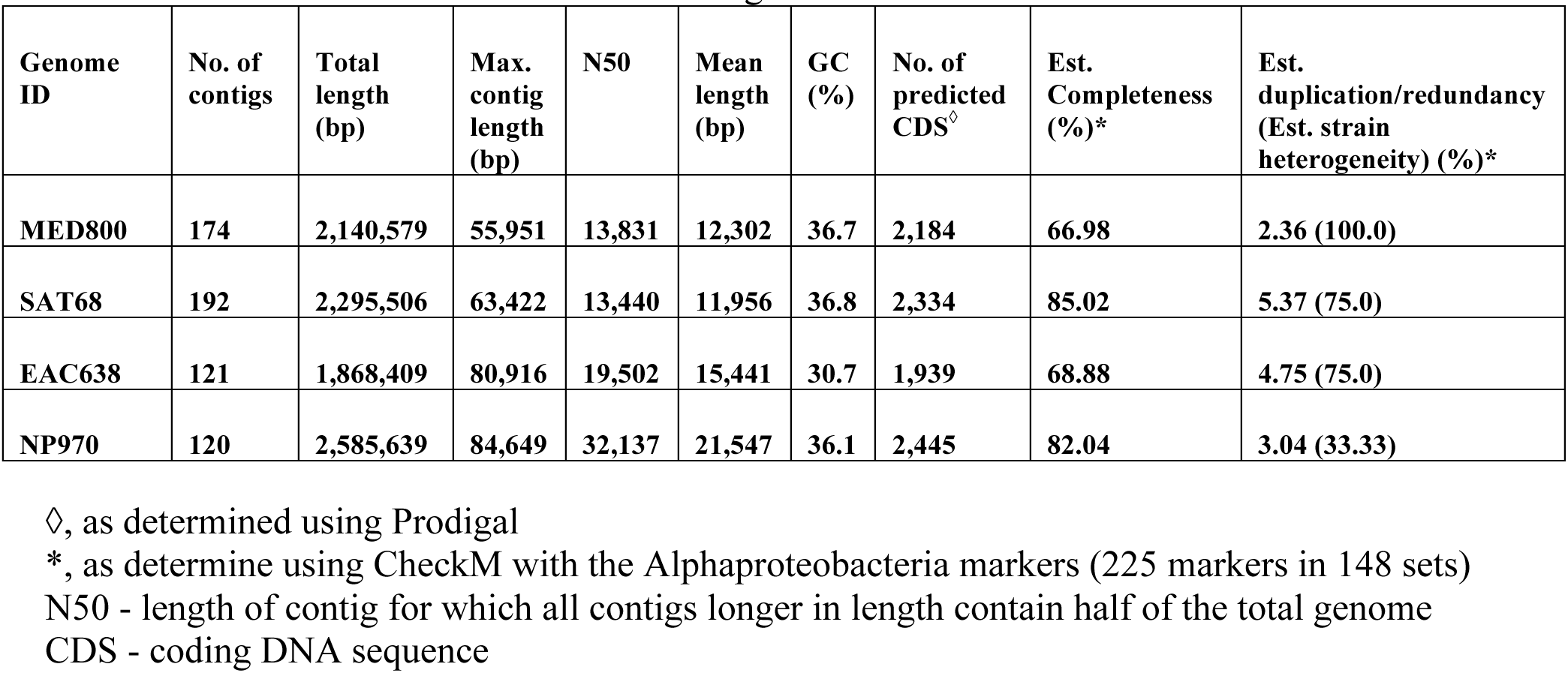
Statistics of the four *Tara* assembled genomes.

**Fig. 4.**
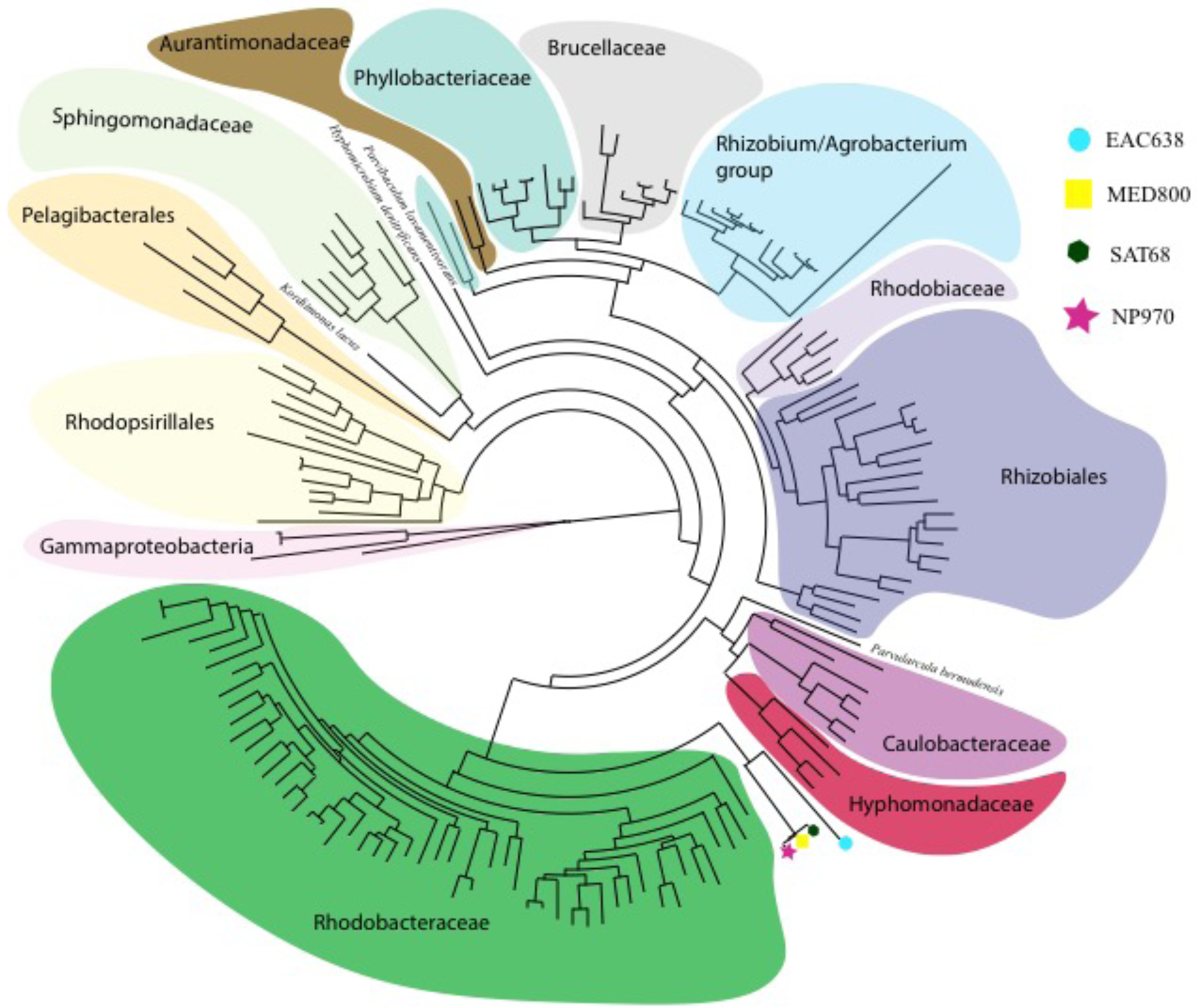
Approximate maximum likelihood *Alphaproteobacteria* phylogenetic tree of 17 concatenated single-copy marker genes for the *Tara* assembled and 160 reference genomes. Reference sequences from the *Gammaproteobacteria* used as an outgroup. Reference genome information, including accession numbers, can be found in Supplementary Data File 4. Phylogenetic distances and local support values can be found in Supplementary Data File 5.

In addition to Rubisco, all four genomes contained genes encoding phosphoribulokinase, an essential gene of the CBB cycle, and 50-89% of the genes necessary to perform complete carbon fixation (Fig. 5). The RCII genes in MED800, EAC638, SAT68, and NP970 were accompanied by bacteriochlorophyll biosynthesis and light-harvesting genes (Supplementary Data File 2). This complement of reaction center, bacteriochlorophyll biosynthesis, and essential carbon fixation genes support a role for autotrophy within these organisms beyond the identified marker genes.

**Fig. 5.**
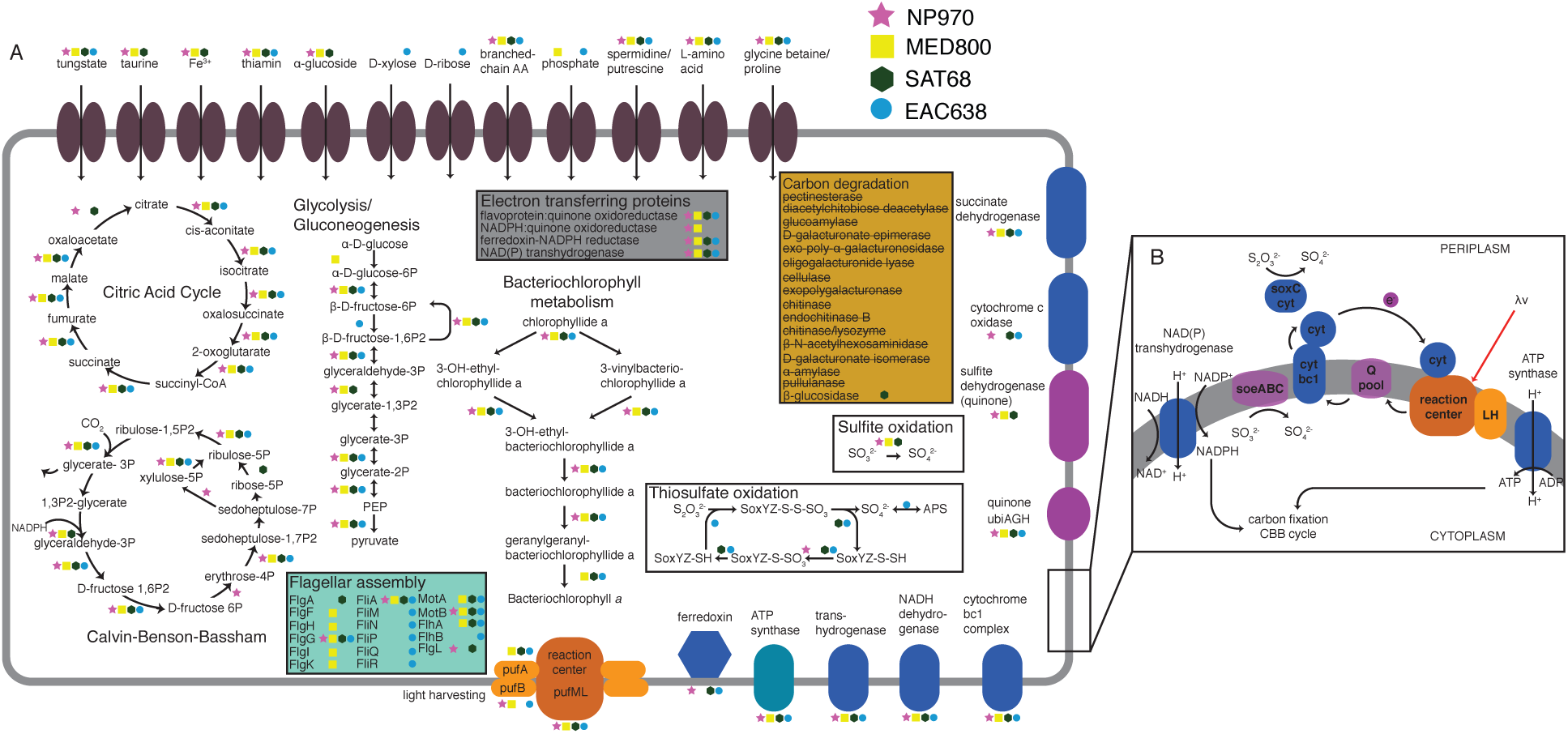
Cellular schematic of the four reconstructed genomes. (**A**) The presence of a gene(s) in a genome is represented by a yellow square (MED800), pink star (NP970), green hexagon (SAT68), and/or blue circle (EAC638). Schematic illustrates predicted membrane bound proteins, but does accurately represent cellular localization. (**B**) A detailed view of the proposed flow of electrons from donors to photosynthesis and carbon fixation. Abbreviations: cyt, cytochrome; Q pool, quinone pool; LH, light-harvesting proteins; soeABC, sulfite dehydrogenase (quinone); CBB, Calvin-Benson-Bassham.

All four genomes possessed ATP-binding cassette (ABC) type transporters for spermidine/putrescine and L- and branched-chain amino acids. These transporters are indicative of the utilization of organic nitrogen compounds, as spermidine and putrescine are nitrogen rich organic compounds, while the scavenging of amino acids reduces the overall nitrogen demands of the cell. Further, the genomes lacked transporters and degradation enzymes for many of the saccharides common in the marine environment^18^ (Fig. 5). However, MED800, SAT68, and NP970 possessed an ABC-type α-glucoside transporter, an annotated β-glucosidase in SAT68, and D-xylose and D-ribose ABC-type transporters in EAC638. While autotrophs are generally considered to not require external sources of organic carbon, saccharide transporters are commonly observed in classical photoautotrophic organisms^19,20^, including the specific example of α-glucoside transporters in strains of *Synechocystis*^21^. For the four genomes, the minimal number of carbon transport and degradation genes may suggest that the organisms have a limited capacity to utilize dissolved organic carbon compounds, but are capable of heterotrophic growth under certain conditions. As such, the genomic potential of these organisms suggest that NP970, SAT68, MED800, and EAC638 are likely facultative autotrophs or mixotrophs.

In oxygenic photosynthesis, electrons are donated as a result of the oxidation of water. Lacking photosystem II, organisms with RCII are incapable of oxidizing water and would require an alternative electron donor for autotrophic processes. EAC638 and SAT68 contained the full/partial suite of genes necessary for thiosulfate oxidation, while MED800, NP970, and SAT68 possessed the genes for oxidizing sulfite. The oxidation of sulfur compounds has previously been linked to autotrophy in the marine environment^22^. The oxidation of organic sulfur compounds, like dimethyl sulfide, has been shown to be a source of thiosulfate^23^ and sulfite^24^ in the marine environment. Electrons derived from inorganic sulfur sources (thiosulfate and/or sulfite) could be transferred directly through cytochromes or membrane-bound quinone dehydrogenases to the electron transport chain. The cycling of electrons through the reaction centers could generate the proton motive force necessary to generate NADH via reverse electron flow^25^, convert NADH to NADPH via transhydrogenase, and generate ATP for the CBB cycle (Fig. 5).

The reconstruction of four genomes from the same novel family in four different provinces (North Pacific, Mediterranean, East Africa coastal current, and South Atlantic) suggests that the observed genomes represent an *in situ* microbial population from the surface marine environment. Each of the genomes recruit metagenomic reads from multiple sampling sites in each province and are present at >0.1% of the microbial relative abundance (range: 0.1-1.04%; mean: 0.286%) in 20 samples (Fig. 1; Supplemental Information 1). Predominantly, the genomes were present in samples are located in the surface (n = 5) or DCM (n = 12). These organisms were collected at depths where light was available for photosynthesis and less frequently identified at deeper depths (n = 3 mesopelagic samples). When >0.1% relative abundance, the genomes tend to be more abundant in the ‘bacterial’/‘girus’ size fraction (n = 14), though were also observed in ‘protistan’ (n = 4), and ‘viral’ (n = 2) size fractions (Supplemental Information 1). The nature of the ‘bacterial’ size fractions suggests that these organisms are generally not particle attached and <1.6μm in size. The occurrence in the protistan fraction may be due to slightly larger cells or attachment to particles, but this data are difficult to interpret as the ‘protistan’ and ‘bacterial’ size fractions can overlap (0.8-1.6μm). As members of the free-living bacterioplankton, these organisms should be poised to grow in aerobic conditions. All four genomes possessed the genes encoding for cytochromes involved in aerobic metabolisms (aa_3_- and bc_1_-type), and lacked the genes for cytochromes involved in microaerobic metabolisms and alternative electron acceptors. Further, all four genomes encoded the gene for an oxygen-dependent ring cyclase (*acsF*), a necessary component in bacteriochlorophyll biosynthesis for which there is alternative that is oxygen-independent (*bchE*) and used by anaerobic organisms.

With this discovery, the potential photosynthesis in the ocean has expanded beyond organisms harboring chlorophyll *a* to include *Alphaproteobacteria* with BChl *a*. Though these organisms have not been cultivated or sequenced before, both PufM and Rubisco in MED800, EAC638, NP970, and SAT68 are phylogenetically related to environmentally-derived protein sequences, lending credence to the fact that these organisms may be a persistent element of oceanic carbon fixation. As such, clades of environmentally sampled genes (*rbsL* and *pufM*) can now be linked to a previously unrecognized source of marine primary productivity. The identification of a globally distributed clade of AAnPs in the ocean capable of carbon fixation continues to expand our understanding of photosynthesis and the marine carbon cycle.

## Materials and Methods

### Assembly

All sequences for the reverse and forward reads from each sampled station and depth within the *Tara Oceans* dataset were accessed from European Molecular Biology Laboratory (EMBL)^10,11^. Typically, *Tara* sampling sites have multiple metagenomic samples, representing different sampling depths and size fractions. The common size fractions were used during sampling were: ‘bacterial’ (0.22-1.6μm) (includes Mediterranean ‘girus’ samples), ‘protistan’ (0.8-5.0μm), ‘girus’ (0.45-0.8μm) and ‘viral’ (<0.22μm). Surface samples were collected at ~5-m depth, while deep chlorophyll maximum (DCM) and mesopelagic depths were variable depending the physiochemical features of the site. Paired-end reads from different filter sizes from each site and depth (e.g., TARA0007, girus filter fraction, sampled at the DCM) were assembled using Megahit^26^ (v1.0.3; parameters: --preset, meta-sensitive) (Supplementary Data File 1). All of the Megahit assemblies from each province were pooled in to two tranches based on assembly size, <2kb and ≥2kb. Longer assemblies (≥2kb) with ≥99% semi-global identity were combined using CD-HIT-EST^27^ (v4.6; -T 90 -M 500000 -c 0.99 -n 10). The reduced set of contiguous DNA fragments (contigs) ≥2kb was then cross-assembled using Minimus2^28^ (AMOS v3.1.0; parameters: -D OVERLAP=100 MINID=95).

### Binning

Contigs from each province were initially clustered into tentative genomic bins using BinSanity^29^. Due to computational limitations, the South Atlantic, East African Coastal province, and Mediterranean Sea were initially run with contig size cutoffs of 11.5kbp, 7.5kbp, and 7kbp, respectively. The BinSanity workflow was run iteratively three times using variable preference values (v.0.2.5.5; parameters: -p [(1) -10, (2) -5, (3) -3] -m 4000 -v 400 -d 0.95). Between each of the three main clustering steps, refinement was performed based on sequence composition (parameters: -p [(1) -25, (2) -10, (3) -3] -m 4000 -v 400 -d 0.95 -kmer 4). After refinement and before the next pass with BinSanity, bins were evaluated using CheckM^30^ (v.1.0.3; parameters: lineage_wf, default settings) for completion and redundancy. Genomes were considered for further analysis based on the completeness and contamination metrics. The cutoff values were: >90% complete with <10% contamination, 80-90% complete with <5% contamination, or 50-80% complete with <2% contamination. Bins meeting these metrics were reclassified as draft genomes were removed from subsequent rounds of clustering. After identification of the four genomes of interest (initially 51-84% complete, <7.0% contamination), binning was performed with CONCOCT^31^ (v.0.4.1; parameters: -c 800 -I 500) on contigs >5kb from each province that had a produced a genome of interest. To improve completion estimates, overlapping CONCOCT and BinSanity bins were visualized using Anvi’o^32^ (v.2.1.0) and manually refined to improve genome completion and minimize contamination estimates (Extended Data Fig. 3-6).

### Annotation

Putative DNA coding sequences (CDSs) were predicted for each genome using Prodigal^33^ (v.2.6.2; -m -p meta). Putative CDS were submitted for annotation by the KEGG database using BlastKOALA^12^ (taxonomy group, Prokaryotes; database, genus_prokaryotes + family_eukaryotes; Accessed March 2017) (Supplementary Data File Table 3). Assessment of pathways and metabolisms of interest were determined using the script KEGG-decoder.py (www.github.com/bjtully/BioData/tree/master/KEGGDecoder). Genomes of interested were determined based on the presence of genes assigned as the M subunit of type-II photochemical reaction center (PufM) and ribulose-1,5-bisphosphate carboxylase (RbsLS). After confirmation of the genes of interest (see below), additional annotations were performed for the genomes using the Rapid Annotation using Subsystem Technology (RAST) service (Classic RAST default parameters - Release70)^34^.

### Phylogeny

An initial assessment of phylogeny was conducted using pplacer^35^ within CheckM. The Prodigal-derived CDSs were searched for a collection of single-copy marker genes that was common to all four *Tara* assembled genomes using hidden Markov models collected from the Pfam database^36^ (Accessed March 2017) and HMMER ^37^ (v3.1b2; parameters: hmmsearch -E1e-10 --notextw). 17 marker genes were identified that met this criteria^38-40^ (Extended Data Table 1). The 17 markers were identified in 2,889 reference genes from complete and partial genomes accessed from NCBI Genbank^41^ (Supplementary Data File 4). If a genome contained multiple copies of a single marker gene both were excluded from the final tree. Only genomes containing ≥10 markers were used for phylogenetic placement. Each marker set was aligned using MUSCLE^42^ (v3.8.31; parameter: -maxiters 8) and trimmed using TrimAL^43^ (v.1.2rev59; parameter: -automated1). Alignments were then manually assessed and concatenated in Geneious^44^. An approximate maximum likelihood tree was generated using FastTree^45^ (v.2.1.10; parameters: -lg -gamma; Supplementary Data File 5). A simplified version of this phylogenetic tree was constructed using the same protocol, but with 160 reference genomes for Fig. 2 (Supplementary Data File 6 and 7).

### Phylogenetic tree – Rubisco and Type-II Reaction Center

RbsL and PufM sequences representing previously described lineages were collected^13,17^ (Supplementary Data File 8 and 9). Additional reference PufM sequences were collected from environmentally generated bacterial artificial chromosomes^5^ and Integrated Microbial Genomes (IMG; Accessed Feb 2017)^46^. Protein sequences from IMG were assessed based on genomes with KEGG Ontology (KO) annotations^47^ for the reaction center subunit M (K08929). PufM sequences from Prodigal predicted CDS (as above) of Global Ocean Survey (GOS) assemblies^16^ were identified using DIAMOND^48^ (v.0.8.36.98; parameters: BLASTP, default settings), where all reference and *Tara* genome sequences were used as a query. Two separate phylogenetic trees were constructed (RbsL and PufM) using the following methodology. Sequences were aligned using MUSCLE^42^ (parameter: -maxiters 8) and automatically trimmed using TrimAL^43^ (parameter: -automated1) (Supplementary Data File 10 and 11). After manual assessment, trimmed alignments were used to construct approximately-maximum-likelihood phylogenetic trees using FastTree^45^ (parameters: -lg -gamma) (Supplementary Data File 12 and 13).

### Relative abundance of genomes in each sample

Reads from each sample were recruited against all assemblies ≥2kb from the same province using Bowtie2^49^ (parameters: default settings), under the assumptions that contigs <2kb would include, low abundance bacteria and archaea, bacteria and archaea with high degrees of repeats/assembly poor regions, fragmented picoeukaryotic genomes, and problematic read sequences (low quality, sequencing artefacts, etc.). For the four sets of contigs (North Pacific, Mediterranean, East Africa Coastal province, and South Atlantic), putative CDS were determined via Prodigal (parameters: see above). In order to estimate the relative abundance of the four analyzed genomes within the bacteria and archaea portion of the total microbial community (excluding eukaryotes and viruses), single-copy marker genes were identified using a collection of previously identified HMMs^50,51^ and searched using HMMER^37^ (hmmsearch -- notextw --cut_tc). Markers belonging to the four genomes were isolated from the total set of environmental markers. The number of reads aligned to each marker was determined using BEDTools^52^ (v2.17.0; multicov default parameters). Length-normalized relative abundance values were determined for each genome as in Equation 1 (Supplementary Data File 1):

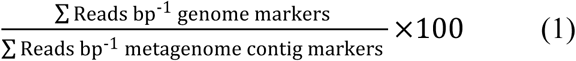

### Data availability

Data is available… submission to NCBI is ongoing. [Currently data is available at FigShare, including high resolution copies of figures, contig and protein sequences, and all supplementary data files: **https://figshare.com/s/9f603e9bbef71164e61b]**

## Acknowledgments

We would like to acknowledge and thank Drs. Eric Webb and William Nelson for providing invaluable comments and critiques in the early stages of this research. We are indebted to the *Tara Oceans* consortium for their commitment to open-access data that allows data aficionados to indulge in the data and attempt to add to the body of science contained within. And we thank the Center for Dark Energy Biosphere Investigations (C-DEBI) for providing funding to BJT and JFH (OCE-0939654). This is C-DEBI contribution number ###.

## Author contributions

BJT conceived of the research plan, performed analysis, and wrote the manuscript. EDG performed analysis and wrote the manuscript. JFH provided funding, provided guidance, and edited the manuscript.

